# Loss of brain insulin production impairs learning and memory in female mice

**DOI:** 10.1101/2025.08.15.670639

**Authors:** Stella K. Baehring, Timothy P. O’Leary, Haoning Howard Cen, Danae Holenka, Kyungchan Kim, Sunday Ajayi, Daniel Semenov, Manuel Belmadani, Hong Li, Arya E. Mehran, Honey Modi, Melissa M. Page, Søs Skovsø, Paul Pavlidis, Shernaz X. Bamji, Eun-Kyoung Kim, James D. Johnson

## Abstract

Diabetes is characterized by dysfunctional insulin release and action, and it is a risk factor for Alzheimer’s disease, the most common form of dementia. Alterations in brain insulin signalling and metabolism have been linked with Alzheimer’s disease. The ancestral insulin gene, *Ins2*, is transcribed locally within the brain. Here we demonstrate that hippocampal *Ins2* mRNA is higher in females than males, and modulated by diet. To specifically determine how locally-produced insulin influences hippocampal function, we used mice with germline *Ins2* knockout (*Ins2*^−/−^) and the normal complement of wildtype *Ins1* alleles. Compensation from the *Ins1* gene ensured normal glucose tolerance, normal insulin sensitivity, normal fasting insulin, and normal body weight under these conditions. Using the Morris water maze, we found that visuo-spatial learning and memory performance of female *Ins2*^−/−^ mice was significantly impaired relative to wild-type mice, whereas the performance of male *Ins2*^−/−^ and wild-type mice did not differ. We profiled isolated hippocampi from female *Ins2*^−/−^ and littermate control mice using RNA sequencing to provide an unbiased analysis of gene expression differences that underlie these behavioural changes. Cyclin D1 (*Ccnd1*) was significantly reduced in *Ins2*^−/−^ mice, prompting us to examine adult neurogenesis using exogenous mitotic marker EdU and the immature neuronal marker doublecortin (DCX). Our data demonstrate female-specific roles for brain-derived *Ins2* on learning and memory function in mice.

**Research in context:** *What is already known about this subject?:*

- Alzheimer’s disease and type 2 diabetes share common risk factors and metabolic dysfunction.
- Insulin and insulin signalling are known to have positive effects on neuronal health and cognition.
- Local insulin production has been reported in the brain, including the hippocampus, which plays critical roles in learning and memory, as well as Alzheimer’s pathophysiology.

*What is the key question?:*

- What are the consequences of deleting the *Ins2* gene, which selectively prevents local brain insulin production, on learning and memory in mice?

*What are the new findings?:*

- Hippocampal insulin production via the *Ins2* gene is sexually dimorphic and modulated by diet.
- Knocking out brain insulin production in *Ins2*^−/−^ mice does not have significant effects on glucose homeostasis in these diet and housing conditions.
- Knocking out brain insulin production in *Ins2*^−/−^ mice results in learning and memory impairments in female but not male mice.

*How might this impact on clinical practice in the foreseeable future?:*

- These pre-clinical observations demonstrate a causal role for insulin in learning and memory, but will require additional human studies before they can impact clinical practice.

## Introduction

Diabetes is a risk factor for dementia [1]. People living with type 2 diabetes can have cognitive impairment similar to early onset Alzheimer’s disease [2]. Alzheimer’s disease often presents with hyperglycemia and insulin dysfunction, including ‘insulin resistance’ in the brain [3–5]. Insulin enhances synaptic plasticity, promotes dendritic spine formation, and increases neurotransmitter turnover [6, 7]. Insulin also influences amyloid clearance and tau phosphorylation [8]. Loss of insulin signaling in astrocytes exacerbates Alzheimer’s pathology [9]. Evidence supporting a role for insulin in cognition in humans and pre-clinical models [10–12] prompted investigators to administer insulin to the brain [13] to treat cognitive dysfunction in neurodegenerative disorders. Indeed, intranasal insulin delivery improves cognitive function in Alzheimer’s disease [10, 13]. Lifestyle interventions that increase insulin sensitivity (e.g. diet, exercise) improve learning and memory, and are also used for Alzheimer’s disease prevention and treatment [7, 14, 15].

Most studies investigating insulin’s effects in the brain are done under the assumption that insulin in the brain is all of pancreatic origin. Although peripheral insulin crosses the blood brain barrier via receptor-mediated transport [16], insulin is also produced locally in the brain [17]. Insulin mRNA has been measured in choroid plexus, cerebral cortex, cerebellum, and hippocampus [18–23]. The specificity of these observations has been confirmed with mice lacking the ancestral insulin 2 (*Ins2*) gene [21, 22], which survive due to the presence of the pancreas-specific insulin coding gene insulin 1 (*Ins1*) [24]. We have shown that *Ins1* does not upregulate its gene expression in the brain to compensate for *Ins2* loss [21, 22]. Insulin synthesis has been reported in hippocampal and olfactory bulb neuronal progenitors [19, 25], sites of adulthood neurogenesis [26, 27]. Insulin stimulates neural stem cell survival, proliferation, and differentiation [28, 29]. These and other data [19–22, 30–33] clearly demonstrate that a small amount of insulin can be synthesized in the brain. Studies that include hippocampal regions consistently report elevated levels of insulin expression relative to other brain regions. The hippocampus is the main region involved in learning and memory, and is one of the first to deteriorate in Alzheimer’s disease. While these lines of evidence point to a role for local *Ins2* production in learning and memory [21, 25], whether brain-derived insulin directly influences learning, memory, and neurogenesis has not been tested.

We used *Ins2* knockout mice to further confirm protein insulin synthesis in the hippocampus and tested how the loss of *Ins2* influences learning and memory in both sexes. We determined that *Ins2* expression in the mouse hippocampus is modulated by diet and biological sex. We found that female mice lacking *Ins2* have deficits in learning and memory. These data provide the first loss-of-function evidence that *Ins2* plays a role in hippocampus-dependant behaviour.

## Materials and methods

### Animals

All procedures were approved by the UBC animal care committee according to Canadian Council on Animal Care guidelines. Five-month C57BL/6J mice were use to assess *Ins2* expression. *Ins2* knockout mice [24] were primarily on a C57BL/6J background. *Ins2*^GFP^ knock-in mice have been described [34]. Mice were housed in same-sex groups of 2-4, in cages with a plastic dome and nesting material. In some cases, mice were singly housed due to aggression or barbering, but this did not differ between genotype or sex. Except as indicated, food and water were *ad libitum*. From weaning, mice were fed chow diet (4.68 kcal/g; 25.3% fat calories, 19.8% protein calories, 54.9% carbohydrate calories; #5015 Lab Diets, Richmond, USA) or a high fat diet (5.56 kcal/g; 58.0% fat calories, 16.4% protein calories, 25.5% carbohydrate calories; #D12330 Open-Source Diets, New Brunswick, USA).

### Immunohistochemistry and immunoblotting

Mice were perfused under isoflurane anesthesia with phosphate-buffered saline (PBS) followed by ice-cold 4% paraformaldehyde in PBS. Brains were post-fixed in paraformaldehyde for 4 hrs, then cryoprotected with 30% sucrose in PBS for 48 hours. Brains were embedded in OCT, frozen on dryice and stored at −80°C, before sectioning at 40µm on a cryostat. Immunostaining of dorsal hippocampal sections used rabbit anti–C-peptide (1:300; Cell Signaling Technology, 4593), rabbit anti–β-gal (1:100; Thermo Fisher Scientific, A-11132) and mouse anti-proinsulin (1:100; R&D Systems, Bio-Techne, MAB13361) primary anti-bodies. Donkey anti-rabbit Cy3 (1:200; Jackson ImmunoResearch, 711-165-152) and donkey anti-mouse Alexa488 (1:200; Jackson ImmunoResearch, 715-545-150) were the secondary antibodies. Hoechst 33258 (1ug/ml, Invitrogen, Thermo Fisher Scientific) stained nuclei. Sections were cover-slipped with VECTASHIELD (Vector Laboratories) and imaged using a LSM 800 confocal microscope (Carl Zeiss).

Immunoblots of extracted proinsulin were conducted as published [22, 35]. Whole hippocampi of 5 *Ins2*^+/+^ and 5 *Ins2*^−/−^ mice were lysed and separated on 15% SDS-PAGE and blotted onto polyvinylidene difluoride membranes (Millipore, IPVH00010) for 30 minutes at 16 V in 25 mM Tris base buffer, pH 7.4, with 192 mM glycine, 10% methanol. Membranes were blocked with 5% skim milk for 1 hour and then incubated with proinsulin antibody (1:1000; Cell Signaling Technology, 8138) or GAPDH (1:10,000; Cell Signaling Technology, 2118) at 4°C overnight. After extensive washing in Tris-buffered saline with 0.1% Tween-20, the membranes were incubated with horseradish peroxidase-conjugated anti-mouse (1:3000; Cell Signaling Technology, 7076S) or anti-rabbit secondary antibody (1:10,000; Thermo, NCI1460KR).

### RNA extraction, cDNA synthesis and quantitative polymerase chain reaction

Brain regions were dissected after CO_2_ euthanasia then snap frozen and stored at −80°C. RNA was isolated with Trizol and chloroform followed by Qiagen RNAeasy (cat #74108 Thermo Fisher). cDNA was synthesized using qScript (cat#101414-100, Quanta Bioscience). Taqman real-time PCR (StepOnePlus, Applied Biosystems) used these primers: *Ins2* Taqman reverse 280→259: GAT CTA CAA TGC CAC GCT TCT G, *Ins2* Taqman probe 224→207: CCT GCT CCC GGG CCT CCA. PCR used 2 min at 50°C, 10 min at 95°C, 40 15-sec cycles at 95°C, 1 min at 60°C. Samples were normalized to beta actin (reverse log of delta). Water and cDNA from *Ins2*^−/−^ tissue were negative controls.

### Metabolic characterization

Metabolic testing was done on individual mice at multiple ages. Mice were fasted for 4 hours prior to intra-peritoneal injections of either glucose (4 mg/kg) or insulin (0.75 U/kg, or 1.5 U/kg) in PBS. Glucose was measured using OneTouch glucometers from the tail. For glucose-stimulated insulin release experiments, blood was collected from the saphenous vein following intra-peritoneal injection of glucose (4 mg/kg). Insulin was quantified using an enzyme linked immunosorbent assay (ALPCO, Salem, USA). For body-weight and fasting blood glucose data, some mice provide data at multiple ages. For tolerance tests and glucose stimulated insulin secretion, data is from different cohorts of mice at each age.

### Morris water maze

The Morris water maze was a white circular pool (110 cm diameter) filled with opaque 23°C water to 16 cm. The escape platform (11.5 cm diameter) was 0.5 cm below the surface. The maze was placed in a diffusely lit room with extra-maze cues. Anymaze (Stoelting) was used to record the movement of mice. Mice completed acquisition training over 6 days (4 trials per day). The escape platform was located in the NW quadrant. For each trial, mice were placed into the pool at one of four release locations and were given a maximum of 60 sec to reach the escape platform. Mice remained on the platform for 5-10 seconds before being returned to the holding cage. If mice did not reach the platform in 60 sec, they were guided. Performance of mice was measured using latency to locate the escape platform, distance travelled to reach the escape platform, cumulative search error and swim-speed. Cumulative search error consisted of the total distance from the escape platform, summed across all recorded positions of the mouse (5 hz). A correction factor is applied, where the CSE value expected for the optimal swim path (determined by start location and swim speed) is subtracted from the total CSE. The day following acquisition training, mice completed a 60 sec probe trial without the escape platform to measure memory. The number of times mice crossed over the location of the escape platform (annulus crossings) and respective locations in other quadrants of the pool were recorded. The day following the acquisition probe, mice completed an additional day of training (acquisition re-training) to determine the extent of extinction from the probe trial, and to re-establish baseline levels of learning performance. Mice then completed reversal training to measure behavioral flexibility and the learning of a new spatial location of the escape platform. During reversal training the escape platform was moved to the opposite side of the pool. Following reversal training, mice completed a 60-sec probe trial to measure memory for the new escape platform location (SE quadrant).

### Anxiety-like behaviour

The same mice tested on the Morris water maze (3 months) also completed testing for anxiety-like behaviour (4-5 months). Movements were recorded with the Ethovision (Noldus) video tracking system. Species typical behaviors were measured with the Boris (Friad and Gamba, 2016) event scoring software. The open field consisted of a box (70 x 70 cm) made of transparent Plexiglas, with 23 cm high walls. For each trial, mice were placed into the one of four corners of the open field (pseudo-randomly determined). Locomotor activity was measured with distance travelled, while anxiety-like behaviour was assessed with entries into the center of the open field (9^th^ of area). Species typical behaviors were also recorded including number of rears, grooming duration, and freezing duration. The light-dark box was constructed from Plexiglas and was divided into two chambers that were either brightly (light zone) or dimly lit (dark zone). A single light placed above the light zone provided lighting in the light-dark box and testing room. Mice were placed in the light zone facing the opening to the dark zone, and the same behavioural measures were recorded in the light-dark box, except that time in the light chamber was used as a measure of anxiety-like behaviour. The elevated plus maze consisted of two pairs of arms, each attached to a central square. One pair of arms had opaque walls around the edge (closed arms), while the other pair of arms had no walls (open arms). A single light provided dim lighting for the elevated plus maze and testing room. Mice were placed onto the central hub of the elevated plus maze, facing a closed arm, and the same behavioral measurements were recorded except that time in the open arms was used as measure of anxiety-like behaviour. The number of times mice dipped their heads over the edge of the open arms (head-dips) was also recorded as a species typical measure of anxiety-like behaviour. Mice same mice completed all three tests, with a 1-week interval between tests.

### Bulk hippocampal RNA sequencing

Sample quality control was performed using the Agilent 2100 Bioanalyzer. Qualifying samples were then prepped following the standard protocol for the NEBnext Ultra ii Stranded mRNA (New England Biolabs). Sequencing was performed on the Illumina NextSeq 500 with Paired End 43bp × 43bp reads. Sequencing data was demultiplexed using Illumina’s bcl2fastq2. De-multiplexed read sequences were then aligned to the *Mus musculous* (PAR-masked)/mm10 reference sequence using STAR aligner. Quality control led to one outlier sample (genotype *Ins2*^−/−^) being removed. Hierarchical clustering and principal component analysis (PCA) identified 3 additional outliers (1 *Ins2*^+/+^, 2 *Ins2*^−/−^) that were removed. Genes with expression below 0.5 counts per million were dropped from the analysis. Differential expression analysis associated with genotype was performed using limma-trend [36] after quantile normalization or using DESeq2 [37], which had consistent results. Pathways were analyzed with clusterProfiler R using Gene Ontology (Biological Process) (FDR < 0.0001) [38] and simplify() function removed redundant GO terms (https://guangchuangyu.github.io/2015/10/use-simplify-to-remove-redundancy-of-enriched-go-terms/). Gene ratio was calculated as the number of leading-edge genes driving the enrichment divided by the total number of detected genes in each gene set. The RNAseq raw data is available at GEO accession GSE305902.

### Quantification of adult hippocampal neurogenesis

Twenty-month-old *Ins2*^+/+^ and *Ins2*^−/−^ mice were given EdU in drinking water (0.25 mg/ml) for 4 weeks. Body weight and water consumption were recorded every week. Mice were transcardially perfused with PBS (0.1M) followed by 4% paraformaldehyde. Brains were extracted, post-fixed in 4% paraformaldehyde overnight, and then transferred to a 30% sucrose solution. Brains were sliced into 40 µm coronal sections using a cryostat. EdU was detected with the Click-iT Plus EdU Alexa Fluor 647 Imaging Kit (Thermo Fisher Scientific). DCX was detected using a rabbit anti-DCX IgG polyclonal antibody (cat# 326 003 Thermo Scientific) and a donkey anti-rabbit IgG H+L Alexa Fluor 488 secondary antibody (cat# A-21206 Invitrogen). We used sections of intestinal villi from EdU treated animals as positive controls for EdU labelling. Stained sections were imaged with a Zeiss LSM 880 confocal microscope (20X objective). Image stacks were stitched and maximum intensity projections were used to count EdU and DCX-positive cells. Six sections were analyzed per animal. The experimenter was blind to conditions.

### Statistical Analysis

For *Ins2* expression in C57BL/6J mice, data were analyzed with sex as a factor when possible. *Ins2*^+/+^ and *Ins2*^−/−^ mice were analyzed separately for each sex and age. For tolerance tests and glucose stimulated insulin secretion, mixed design ANOVAs (2 x 6 or 2 x 3, respectively) were used with genotype as a between subject factor and sampling time as a within subject factor. For acquisition and reversal training in the MWM, analyses were completed separately for days 1-3 and days 4-6, which reflect initial learning and near-asymptotic performance respectively. Measures of learning performance were analyzed with mixed design ANOVAs (2 x 3), using genotype as a between subject factor and day of training as within subject factor. All other analyses were completed using one-way between subject ANOVAs comparing genotypes. Post-hoc tests were completed using unpaired t-tests, with Bonferonni corrections applied independently for each statistical test.

## Results

### Ins2 is produced in the brain, expressed more in females, and modulated by diet

We previously reported *Ins2* mRNA and insulin protein in the mouse brain [21], but did not show the specificity of hippocampal C-peptide and proinsulin immunoreactivity. Here, we found C-peptide, a byproduct of insulin synthesis, particularly in dentate gyrus-CA3 (**Figure 1A)**. This was confirmed with immunoblots showing a small but clear *Ins2*-specific band for proinsulin in hippocampus lysates, as well as the expected prominent band in pancreas (**Figure 1B**). No insulin protein was found in hippocampi from *Ins*2^−/−^ because the non-ancestral *Ins1* gene is not robustly expressed in the mouse brain [21]. Proinsulin staining in the CA3 and choroid plexus was also largely absent in *Ins2*^−/−^ mice (**Figure 1C**). Importantly, in *Ins2*^+/−^ mice, proinsulin labelling colocalized with β-gal labelling, a proxy for *Ins2* gene activity. We also visualized green fluorescence in cultured hippocampal neurons from mice with GFP knocked into the endogenous *Ins2* locus (**Figure 1D**), which we have previously shown correlates with *Ins2* mRNA, pre-mRNA, and proteins levels in the pancreas [34]. We quantified *Ins2* mRNA across 6 brain regions in C57BL/6J mice. Brain *Ins2* expression was statistically different between regions (F(5,70) = 18.42, p< 0.0001), being highest in hippocampus and cerebellum (**Figure 2A**). We observed sex differences (interaction, (F(5,70) = 2.67, p < 0.05), with; females showing higher expression of *Ins2* than males in the hippocampus and cerebellum (post-hoc, p < 0.005) (**Figure 2A**). Because pancreatic *Ins2* is modulated by diet, we tested whether similar regulation occurs in brain. In female mice, *Ins2* expression was reduced after high-fat diet when compared to a control diet (**Figure 2B,C**). These data demonstrate that *Ins2* mRNA levels differ between brain regions, sexes, and dietary contexts. We examined possible effects of age on brain *Ins2* mRNA in recent publicly accessible data [39], but we did not find significant differences.

**Figure 1.**
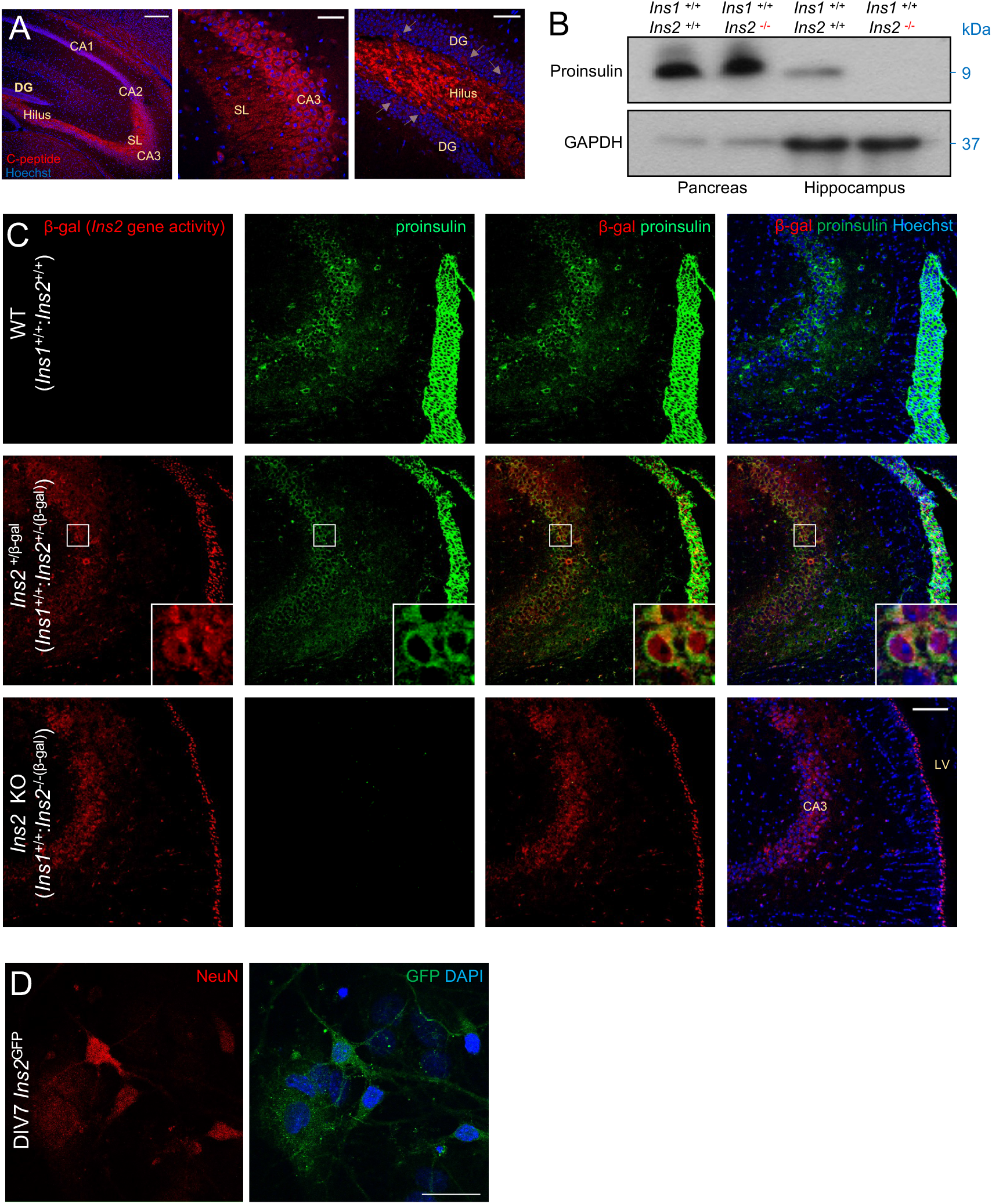
Specific *Ins2* protein immunoreactivity in mouse brain. **(A)** Immunoreactivity for C-peptide, a product of insulin biosynthesis, is detected in the hilus, stratum lucidum (SL), and CA3 regions of hippocampus. C-peptide immunoreactive cells were observed in the granule cell layer of the dentate gyrus, as indicated by arrows. Scale bar, 100 μm. **(B)** Immunoblot of proinsulin expression in the pancreas and hippocampus from *Ins2*^+/+^ and *Ins2*^−/−^ mice. **(C)** Proinsulin-positive cells were detected and co-localized with β-gal in the CA3 regions. β-galactosidase (β-gal) staining was detected in *Ins2* knockout and heterozygous mice but not wildtype (WT). Pro-insulin was detected in WT and β-gal heterozygous mice but not knockout mice. Inset box, 20 μm. **(D)** GFP fluorescence in hippocampal neuron cultures of from *Ins2*^GFP/GFP^ knock-in, replacement mice. Scale bar, 10 μm. For these qualitative data, each analysis was repeated at least 3 times. Mice were 2-5 months of age.

**Figure 2.**
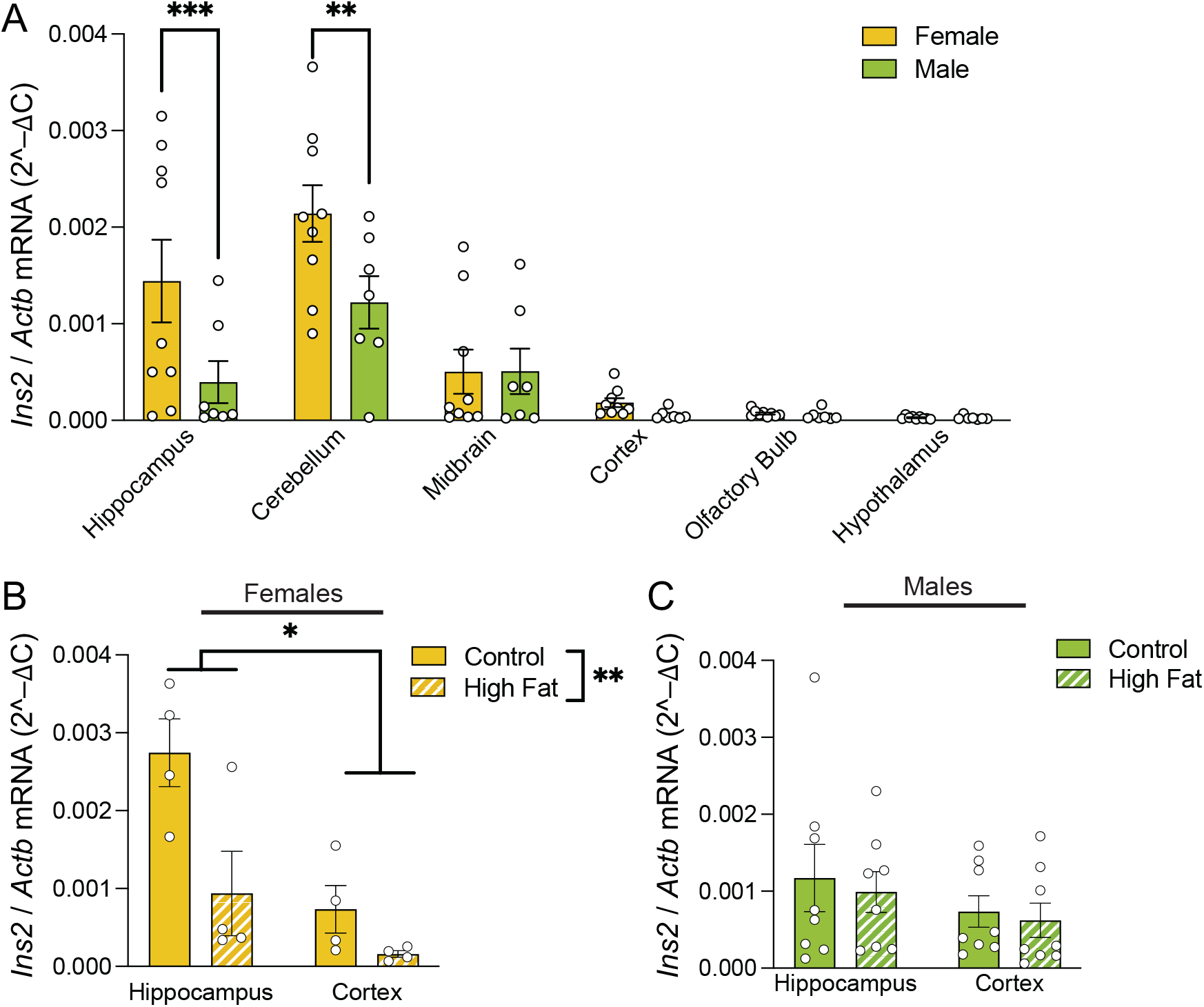
*Ins2* expression in the brain is sexually dimorphic and is modulated by diet in female mice. **A)** *Ins2* mRNA in male and female mice across multiple brain regions, including the hippocampus, cerebellum, cerebral cortex, olfactory bulb, hypothalamus and midbrain. *Ins2* mRNA was normalized to beta-actin mRNA. **(B,C)** *Ins2* mRNA in the hippocampus and cerebral cortex of male and female mice fed high-fat or control chow diets. Mice were 5 months of age. * = p < 0.05; ** = p < 0.005; *** = p < 0.0001.

### Ins1 is sufficient to preserve normal peripheral metabolism in the absence of Ins2

Before using *Ins2*^−/−^ mice to examine the roles of *Ins2* in the brain, it was critical to confirm whole body metabolism was not significantly affected. Previous studies have suggested that *Ins1* can compensate for loss of *Ins2* [21, 24, 40]. However, previous work only studied males so we characterized metabolism from 3-16 months of age in littermate male and female *Ins2*^−/−^ mice side-by-side. Body weight did not differ between *Ins2*^−/−^ and *Ins2*^+/+^ mice at any age **(Figure 3A,B)**. Fasted blood glucose was similar between *Ins2*^−/−^ and *Ins2*^+/+^ mice (**Figure 3C,D)**. *Ins2*^−/−^ and *Ins2*^+/+^ mice did not differ in glucose tolerance or insulin tolerance tests **(Figure 3E-L)**. *Ins2*^−/−^ and *Ins2*^+/+^ mice showed similar glucose stimulated insulin secretion (**Figure 3M-P**), although there was a trend for reduced insulin in female *Ins2*^−/−^ mice compared with *Ins2*^+/+^ mice at the oldest age measured (**Figure 3N**). These data suggest that *Ins1* can compensate for *Ins2* knockout under these diet and housing conditions.

**Figure 3.**
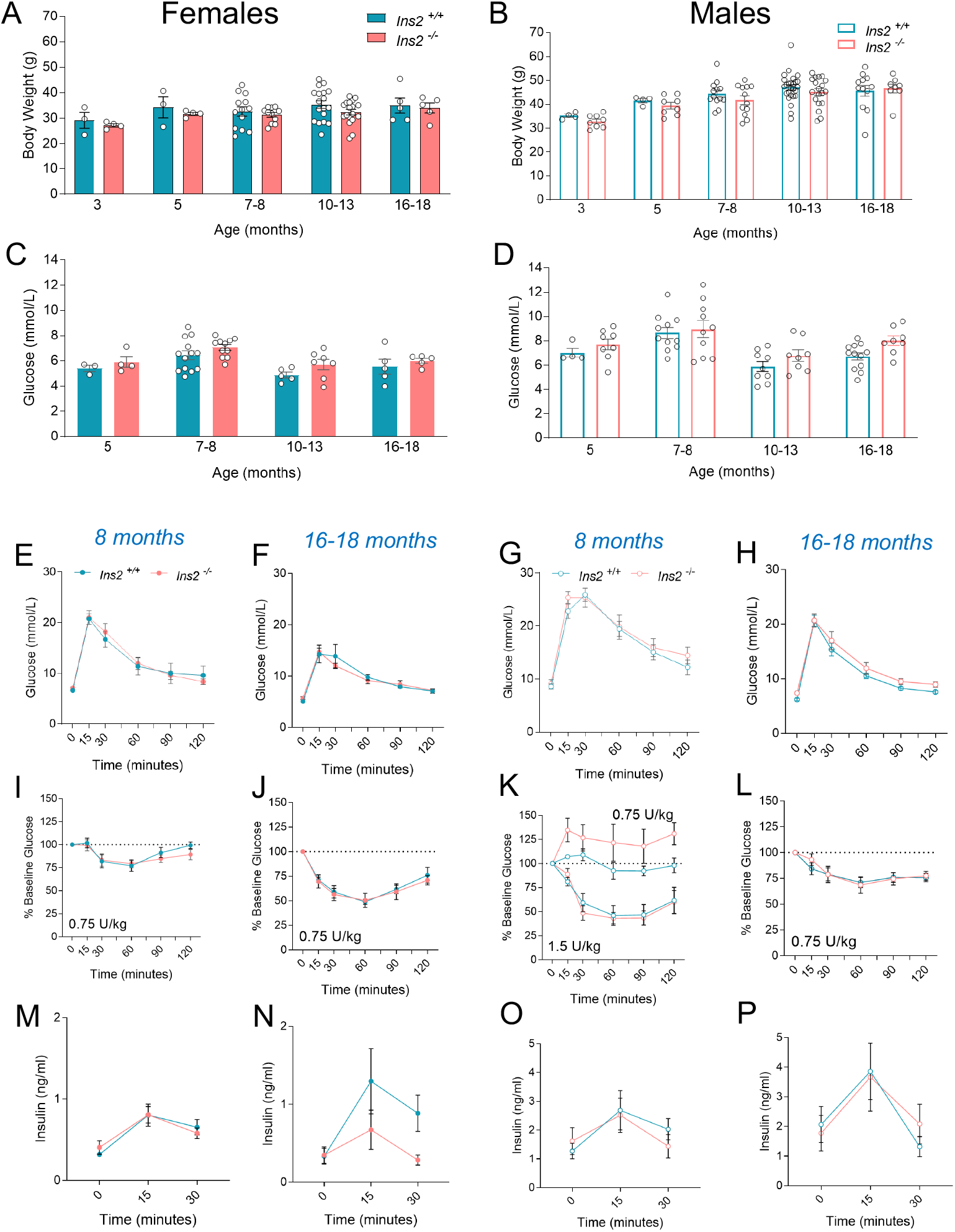
Metabolic profile of young and old *Ins2*^−/−^ mice on a chow diet. **(A,B)** *Ins2*^−/−^ and *Ins2*^+/+^ mice did not differ in body weight. **(C,D)** *Ins2*^−/−^ and *Ins2*^+/+^ mice did not differ in fasted blood glucose. **(E-L)** *Ins2*^−/−^ and *Ins2*^+/+^ mice did not differ in either glucose tolerance or insulin tolerance tests. **(M-P)** *Ins2*^−/−^ and *Ins2*^+/+^ mice were also similar in glucose stimulated insulin secretion, although there was a trend for higher insulin in female *Ins2*^+/+^ than *Ins2*^−/−^ mice in old age (16-18 months). For E-P, 4-13 mice per group.

### Ins2 loss impairs visuo-spatial learning and memory in female, but not male mice

Given *Ins2* expression is prominent in the hippocampus relative to other neuronal populations, we examined whether hippocampal-dependent behaviour was affected by *Ins2*^−/−^ ablation in the Morris water maze [41]. Because we observed sexual dimorphism in hippocampal *Ins2* mRNA, we compared both females (**Figure 4A-F**) and males (**Figure 4G-L**) at 3 months of age. During acquisition training, female *Ins2*^−/−^ and *Ins2*^*+/+*^ mice did not differ in latency (**Figure 4A**), distance (**Figure 4D**), or cumulative search error (**Figure 4B**). During the acquisition probe memory test, however, female *Ins2*^−/−^ completed fewer crossings over the location of the escape platform (annulus crossings) than *Ins2*^*+/+*^ mice **(Figure 4C)**. We provided an additional day of acquisition training (re-training, RT) to reduce potential extinction effects, and to provide another indirect measure of memory (assessing retention of learning performance from 2 days prior). On the re-training day, female *Ins2*^−/−^ performed worse than *Ins2*^*+/+*^ mice on all measures of learning (**Figure 4A,B,D**). During the first 3 days of reversal training, female *Ins2*^−/−^ and *Ins2*^+/+^ did not differ on measures of learning; over the last 3 days of reversal training, however, female *Ins2*^−/−^ mice performed worse than *Ins2*^*+/+*^ mice on all measures of learning (**Figure 4A,B,D**). During the reversal probe memory test, *Ins2*^−/−^ mice completed fewer crossings over the escape platform location than *Ins2*^*+/+*^ mice **(Figure 4D)**. We measured swim speed throughout acquisition and reversal training, which can reflect motivation to find the platform. Female *Ins2*^−/−^ mice swam quicker than *Ins2*^*+/+*^ mice during the first 3 days of acquisition training, but this difference was not present during the last 3 days of acquisition training, nor at any point during reversal training **(Figure 4E)**. We conclude that differences in swim-speed do not account for the impaired learning and memory observed in *Ins2*^−/−^ mice. These results indicate impaired learning and memory in *Ins2*^−/−^ mice. We assessed learning and memory in male *Ins2*^−/−^ and *Ins2*^*+/+*^ mice in the Morris water maze but did not find any differences in learning performance **(Figure 4G,H,J,K,I,L)**. Unlike females, loss of *Ins2*^−/−^ in male mice is not sufficient to disrupt visuo-spatial learning or memory.

**Figure 4.**
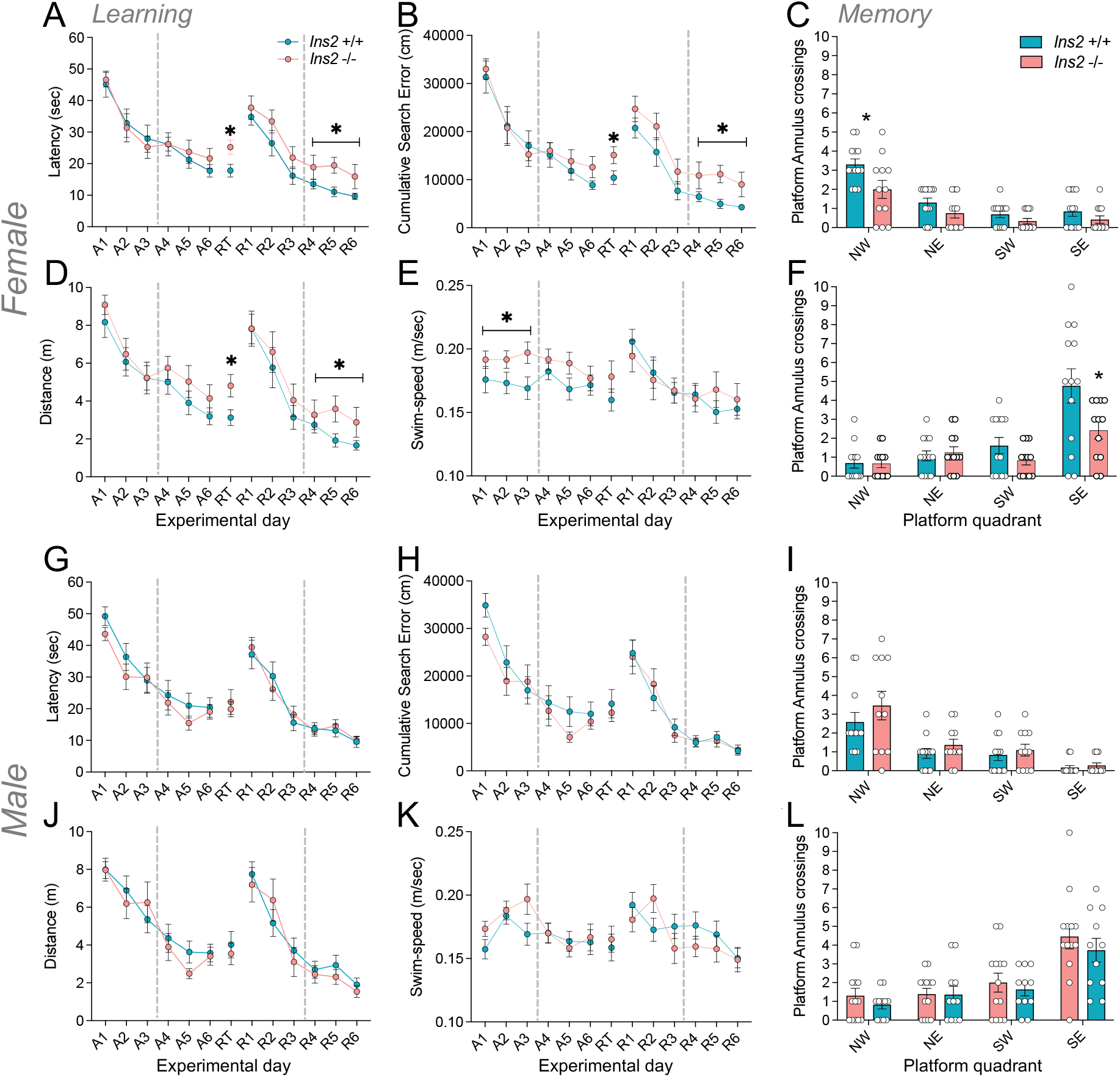
Visuo-spatial learning and memory is impaired in female *Ins2*^−/−^ mice. **(A)** Latency to locate the escape platform in female *Ins2*^−/−^ mice and *Ins2*^+/+^ mice during acquisition (A1-6), re-training (RT) and reversal training (R1-6). Performance for the first 3 days and last 3 days of acquisition and reversal training were analyzed separately (indicated by horizontal dashed line). **(B)** Cumulative search error for female *Ins2*^−/−^ and *Ins2*^*+/+*^ mice on the same study days. **(C)** Platform annulus crossings completed by female *Ins2*^−/−^ and *Ins2*^+/+^ mice during the acquisition probe trial. **(D)** Distance travelled by female *Ins2*^−/−^ and *Ins2*^*+/+*^ mice to reach the escape platform on each study day. **(E)** Swim-speed for female *Ins2*^−/−^ and *Ins2*^*+/+*^ mice over acquisition and reversal training. **(F)** Platform annulus crossings during the reversal probe trial. **(G-L)** Show the same measurements as above, but for male mice. Mice were 3 months of age, 11-13 mice per group * = p < 0.05

### Loss of Ins2 does not influence anxiety-like behaviour or locomotor activity

The dorsal hippocampus is involved in visuo-spatial navigation and memory, while the ventral hippocampus is involved moreso in anxiety-like behaviour [42, 43]. We examined anxiety-like behaviour in 4-5 month *Ins2*^−/−^ mice using three common tests of anxiety-like behaviour: the open-field, light-dark box, and elevated-plus maze. All tests produced marked avoidance of anxiogenic areas, with mice spending less time in anxiogenic regions than expected by chance **(ESM Figure 1)**. For both males and females, however, *Ins2*^−/−^ and *Ins2*^+/+^ mice did not differ in anxiety-related behaviour. We also extended our analysis to include locomotor activity and species-typical measures of exploration and anxiety-like behaviour (i.e. grooming, rearing, freezing, etc.), but did not find any differences between *Ins2*^−/−^ and *Ins2*^+/+^ mice **(ESM Figure 1)**. Thus, the effects of *Ins2* knockout in female mice appear to be specific to learning and memory in the Morris water maze.

### Transcriptomics of hippocampi from aged female Ins2^−/−^ mice

We conducted bulk hippocampal RNAseq in female *Ins2*^−/−^ mice (12 months) to identify molecular mechanisms that could underlie the defects in learning and memory. We started with 11 samples in each group, but 3 samples (6, 12 and 17) were removed because they were outliers based on hierarchical clustering and PCA (**Figure 5A,B**). *Ins2* mRNA was absent in *Ins2*^−/−^ samples **(Figure 5C**). We identified 6 downregulated and 13 upregulated genes in *Ins2*^−/−^ *mice* (**Figure 5D**). The cell cycle regulatory gene *Ccnd1* and the stem cell regulatory long non-coding RNA Gm26793 [44] were the most significantly downregulated and upregulated genes, respectively. These genes were robustly altered even when the outlier samples were included. Gene set enrichment analysis showed that the (semi-redundant) gene ontology terms “aerobic respiration” and “oxidative phosphorylation” were the most significantly upregulated (**Figure 5E**). Insulin regulates oxidative phosphorylation in many tissues, including brain [45]. The gene ontology term “epithelium development” was the only significantly downregulated pathway (**Figure 5E**). These relatively well-powered transcriptomic studies point to potential molecular mechanisms underlying the effects of insulin signalling the in female rodent hippocampus.

**Figure 5.**
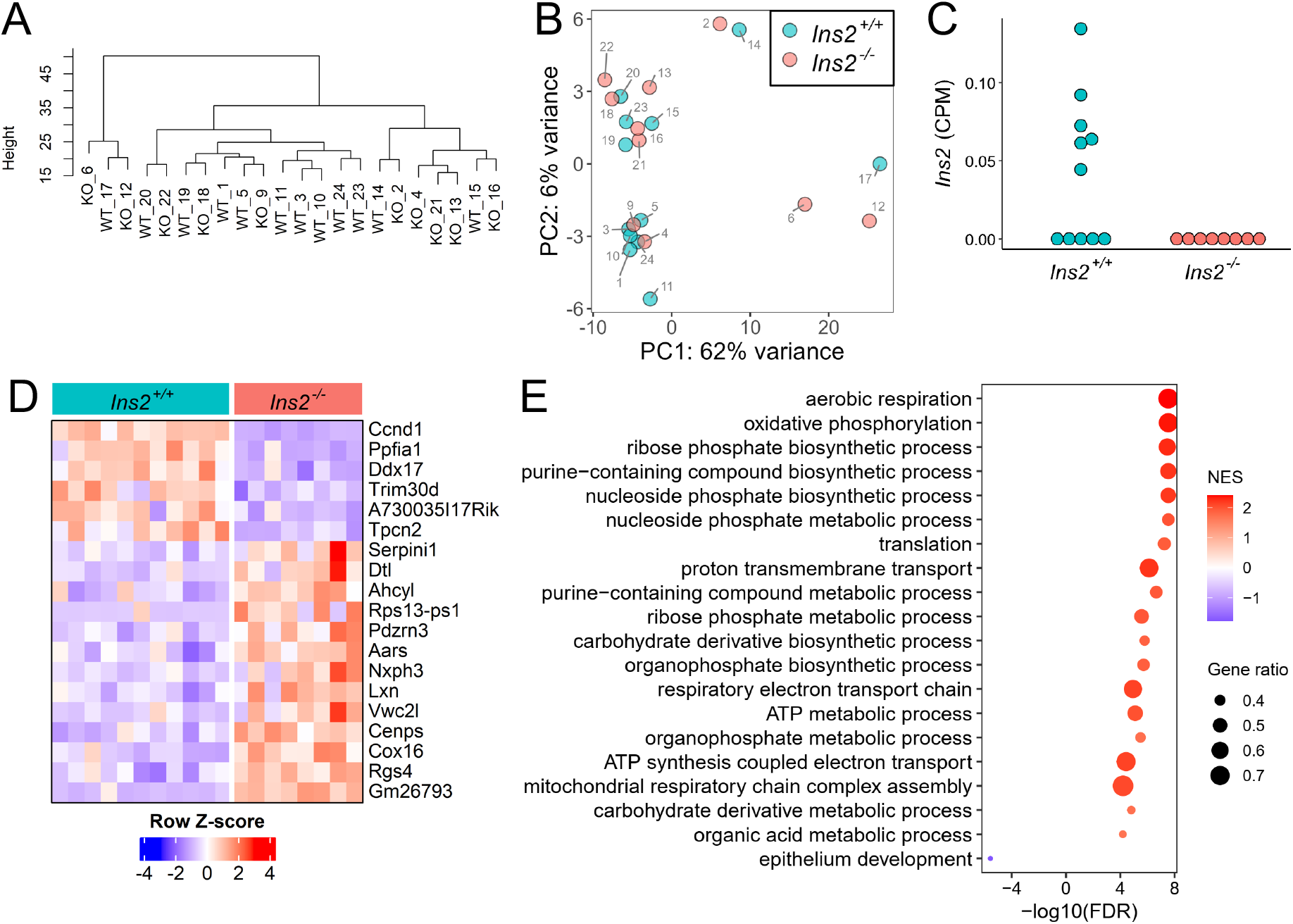
Bulk mRNA sequencing and cluster analysis of isolated *Ins2*^−/−^ and *Ins2*^+/+^ hippocampi from female mice. **(A)** Cluster dendrogram reflects a measure of the distance between samples (height) and **(B)** Principal component analysis showing the 3 outlier samples (6, 12, 17). **(C)** No *Ins2* mRNA was detected in *Ins2*^−/−^ hippocampi. **(D)** Differentially expressed genes ranked from most significantly downregulated to upregulated (top to bottom). **(E)** Gene set enrichment analysis using Gene Ontology - Biological Process shows the normalized enrichment scores (NES; FDR< 0.0001).

### Neurogenesis in Ins2^−/−^ mice

Given that *Ins2*^−/−^ mice had reduced *Ccnd1* mRNA, a gene with roles in neurogenesis, we asked if hippocampal neurogenesis was altered in aged *Ins2*^−/−^ mice (20 months). We first confirmed the robustness of our staining by confirming Edu+ cells in intestinal villi, a region with high cell proliferation (**ESM Figure 2A)**. In female brains, the densities of Edu+ cells and DCX+ cells did not differ between *Ins2*^−/−^ and *Ins2*^+/+^ mice (**ESM Figure 2B-F**). In male brains, *Ins2*^−/−^ mice had a larger density of EdU+ cells in the dorsal (F(1,10) = 5.57, p<0.05)(**ESM Figure 2C**), but not ventral hippocampus (**ESM Figure 2D**). We have shown that reducing *Ins2* has cell-autonomous proliferative effects in the pancreas [46]. One could speculate this is compensation preventing male *Ins2*^−/−^ mice from developing deficits in learning and memory. The density of DCX+ cells did not differ between *Ins2*^−/−^ and Ins2^+/+^ male mice in either the dorsal or ventral hippocampus (**ESM Figure 2E,F**). These data suggest that adult hippocampal neurogenesis, measured at this stage of life, is not a major factor in differences in learning and memory in female *Ins2*^−/−^ mice.

## Discussion

The objectives of this study were to further elucidate the expression of insulin in the hippocampus and to determine the role of brain-produced insulin on hippocampus-dependent learning and memory, loss-of-function causality studies that cannot be conducted in humans. Given that *Ins2* is expressed in both the brain and pancreas, it was important to confirm that *Ins2* knockout did not alter peripheral metabolism, which could theoretically affect brain function indirectly. Using mice devoid of hippocampal pro-insulin we defined a female-specific role for brain-insulin production in learning and memory. To the best of our knowledge, this is the first time mammalian cognitive behaviour has been examined in the context of genetic insulin reduction. We found sexual dimorphic *Ins2* expression and environmental modulation in the hippocampus, as well as female-specific consequences of *Ins2* loss. Women are at higher risk to develop early onset Alzheimer’s disease [47]. The female-specific memory impairment observed after loss of *Ins2*, highlights the importance of including females in pre-clinical research with insulin and cognitive dysfunction, and predicts that insulin administration may be more effective at treating cognitive impairment in females than males. Importantly, when compared to other studies, our RNA-seq experiment was conducted on middle-aged mice, which is especially relevant to understanding the potential role of *Ins2* in the development of mild-cognitive impairment and early onset Alzheimer’s disease.

Previous studies have ablated insulin-expressing neurons with the beta cell toxin streptozotocin [49–51], but such drugs may affect other areas of the body and brain. Although the present study circumvents some of these issues, the *Ins2* knockout model has limitations. While we observed a female-specific effect, we have no information on the estrous status of these mice. With a global *Ins2* knockout, it cannot be entirely excluded that the results of this study are exclusively due to brain-insulin loss. Generating tissue-specific *Ins2* knockout could circumvent this limitation, but our efforts to do so have been unsuccessful. In addition, the loss of *Ins2* was lifelong, and thus effects of *Ins2* loss during early development cannot be excluded. Future studies could employ a floxed *Ins2* allele and an inducible brain-specific Cre driver to selectively knock out *Ins2* during adulthood. Another limitation of our work is that the behavioural, RNA-seq, and neurogenesis experiments were conducted at different ages, which may limit our ability to resolve mechanisms for *Ins2* in hippocampal dependant memory. Our work also does not identify the cellular target(s) of local insulin production, nor their molecular mechanisms. Insulin receptors have been shown to mediate important roles of insulin in many neuronal cell populations in the brain, including the hippocampus [52–54]. Insulin also acts directly though insulin receptors to modulate the activity of non-neuronal brain cells including astrocytes [55–57]. Loss of insulin action in astrocytes significantly worsened Alzheimer’s pathologies in a mouse model [9]. Cerebrovascular insulin receptors were also shown to be deficient in Alzheimer’s disease [58]. More research is required to elucidate local insulin-insulin receptor trophic circuits across cell types in the brain, including in humans.

## Supporting information

ESM

## Acknowledgments

We thank many colleagues for helpful discussions. The graphical abstract was created in BioRender (Johnson, J. (2025) https://BioRender.com/awrjqmn)

## References

[1] Hölscher C (2011) Diabetes as a risk factor for Alzheimer’s disease: insulin signalling impairment in the brain as an alternative model of Alzheimer’s disease. In. Portland Press Ltd.

[2] Ryan CM, Geckle M (2000) Why is learning and memory dysfunction in Type 2 diabetes limited to older adults? Diabetes/metabolism research and reviews 16(5): 308–315

[3] De Felice FG, Lourenco MV, Ferreira ST (2014) How does brain insulin resistance develop in Alzheimer’s disease? Alzheimer’s & Dementia 10(1): S26–S32

[4] Talbot K, Wang H-Y, Kazi H, et al. (2012) Demonstrated brain insulin resistance in Alzheimer’s disease patients is associated with IGF-1 resistance, IRS-1 dysregulation, and cognitive decline. Journal of Clinical Investigation 122(4): 1316–1338. 10.1172/jci59903

[5] Ma L, Wang J, Li Y (2015) Insulin resistance and cognitive dysfunction. Clinica chimica acta 444: 18–23

[6] Forner S, Baglietto-Vargas D, Martini AC, Trujillo-Estrada L, LaFerla FM (2017) Synaptic Impairment in Alzheimer’s Disease: A Dysregulated Symphony. Trends Neurosci 40(6): 347–357. 10.1016/j.tins.2017.04.002

[7] Kellar D, Craft S (2020) Brain insulin resistance in Alzheimer’s disease and related disorders: mechanisms and therapeutic approaches. Lancet Neurol 19(9): 758–766. 10.1016/S1474-4422(20)30231-3

[8] Laws SM, Gaskin S, Woodfield A, et al. (2017) Insulin resistance is associated with reductions in specific cognitive domains and increases in CSF tau in cognitively normal adults. Scientific reports 7(1): 1–11

[9] Chen W, Huang Q, Lazdon EK, et al. (2023) Loss of insulin signaling in astrocytes exacerbates Alzheimer-like phenotypes in a 5xFAD mouse model. Proceedings of the National Academy of Sciences 120(21): e2220684120. doi:10.1073/pnas.2220684120

[10] Freiherr J, Hallschmid M, Frey WH, et al. (2013) Intranasal insulin as a treatment for Alzheimer’s disease: a review of basic research and clinical evidence. CNS drugs 27(7): 505–514

[11] Zhang H, Hao Y, Manor B, et al. (2015) Intranasal insulin enhanced resting-state functional connectivity of hippocampal regions in type 2 diabetes. Diabetes 64(3): 1025–1034

[12] Li X, Yang Y, Bai X, et al. (2024) A brain-derived insulin signal encodes protein satiety for nutrient-specific feeding inhibition. Cell Rep 43(6): 114282. 10.1016/j.celrep.2024.114282

[13] Born J, Lange T, Kern W, McGregor GP, Bickel U, Fehm HL (2002) Sniffing neuropeptides: a transnasal approach to the human brain. Nature Neuroscience 5(6): 514–516. 10.1038/nn0602-849

[14] Ko Y, Chye SM (2020) Lifestyle intervention to prevent Alzheimer’s disease. Reviews in the Neurosciences 31(8): 817–824

[15] Braak H, Braak E (1995) Staging of Alzheimer’s disease-related neurofibrillary changes. Neurobiology of aging 16(3): 271–278

[16] Gray SM, Aylor KW, Barrett EJ (2017) Unravelling the regulation of insulin transport across the brain endothelial cell. Diabetologia 60(8): 1512–1521. 10.1007/s00125-017-4285-4

[17] Dakic T, Jevdjovic T, Lakic I, et al. (2023) The Expression of Insulin in the Central Nervous System: What Have We Learned So Far? International Journal of Molecular Sciences 24(7): 6586

[18] Molnar G, Farago N, Kocsis AK, et al. (2014) GABAergic neurogliaform cells represent local sources of insulin in the cerebral cortex. J Neurosci 34(4): 1133–1137. 10.1523/JNEUROSCI.4082-13.2014

[19] Nemoto T, Toyoshima-Aoyama F, Yanagita T, et al. (2014) New insights concerning insulin synthesis and its secretion in rat hippocampus and cerebral cortex: amyloid-beta1-42-induced reduction of proinsulin level via glycogen synthase kinase-3beta. Cell Signal 26(2): 253–259. 10.1016/j.cellsig.2013.11.017

[20] Mazucanti CH, Liu QR, Lang D, et al. (2019) Release of insulin produced by the choroid plexis is regulated by serotonergic signaling. JCI Insight 4(23). 10.1172/jci.insight.131682

[21] Mehran AE, Templeman NM, Brigidi GS, et al. (2012) Hyperinsulinemia drives diet-induced obesity independently of brain insulin production. Cell Metab 16(6): 723–737. 10.1016/j.cmet.2012.10.019

[22] Lee J, Kim K, Cho JH, et al. (2020) Insulin synthesized in the paraventricular nucleus of the hypothalamus regulates pituitary growth hormone production. JCI Insight 5(16). 10.1172/jci.insight.135412

[23] Mazucanti CH, Kennedy V, Jr., Premathilake HU, et al. (2023) AAV5-mediated manipulation of insulin expression in choroid plexus has long-term metabolic and behavioral consequences. Cell Rep 42(8): 112903. 10.1016/j.celrep.2023.112903

[24] Duvillie B, Cordonnier N, Deltour L, et al. (1997) Phenotypic alterations in insulin-deficient mutant mice. Proc Natl Acad Sci U S A 94(10): 5137–5140

[25] Kuwabara T, Kagalwala MN, Onuma Y, et al. (2011) Insulin biosynthesis in neuronal progenitors derived from adult hippocampus and the olfactory bulb. EMBO Mol Med 3(12): 742–754. 10.1002/emmm.201100177

[26] Berdugo-Vega G, Dhingra S, Calegari F (2023) Sharpening the blades of the dentate gyrus: how adult-born neurons differentially modulate diverse aspects of hippocampal learning and memory. Embo J 42(22): e113524. 10.15252/embj.2023113524

[27] Hitoshi S, Mandyam CD, Shetty AK (2023) Editorial: Advances in adult neurogenesis. Front Neurosci 17: 1307844. 10.3389/fnins.2023.1307844

[28] Brooker GJ, Kalloniatis M, Russo VC, Murphy M, Werther GA, Bartlett PF (2000) Endogenous IGF-1 regulates the neuronal differentiation of adult stem cells. Journal of neuroscience research 59(3): 332–341

[29] Åberg MA, Åberg ND, Palmer TD, et al. (2003) IGF-I has a direct proliferative effect in adult hippocampal progenitor cells. Molecular and Cellular Neuroscience 24(1): 23–40

[30] Young III WS (1986) Periventricular hypothalamic cells in the rat brain contain insulin mRNA. Neuropeptides 8(2): 93–97

[31] Havrankova J, Schmechel D, Roth J, Brownstein M (1978) Identification of insulin in rat brain. Proceedings of the National Academy of Sciences 75(11): 5737–5741

[32] Csajbok EA, Tamas G (2016) Cerebral cortex: a target and source of insulin? Diabetologia 59(8): 1609–1615. 10.1007/s00125-016-3996-2

[33] Kuwabara T, Kagalwala MN, Onuma Y, et al. (2011) Insulin biosynthesis in neuronal progenitors derived from adult hippocampus and the olfactory bulb. EMBO Molecular Medicine 3(12): 742–754. 10.1002/emmm.201100177

[34] Chu CMJ, Modi H, Ellis C, et al. (2022) Dynamic Ins2 Gene Activity Defines beta-Cell Maturity States. Diabetes 71(12): 2612–2631. 10.2337/db21-1065

[35] Kim S, Kim N, Park S, et al. (2020) Tanycytic TSPO inhibition induces lipophagy to regulate lipid metabolism and improve energy balance. Autophagy 16(7): 1200–1220. 10.1080/15548627.2019.1659616

[36] Ritchie ME, Phipson B, Wu D, et al. (2015) limma powers differential expression analyses for RNA-sequencing and microarray studies. Nucleic Acids Research 43(7): e47–e47. 10.1093/nar/gkv007

[37] Love MI, Huber W, Anders S (2014) Moderated estimation of fold change and dispersion for RNA-seq data with DESeq2. Genome Biol 15(12): 550. 10.1186/s13059-014-0550-8

[38] Yu G, Wang LG, Han Y, He QY (2012) clusterProfiler: an R package for comparing biological themes among gene clusters. OMICS 16(5): 284–287. 10.1089/omi.2011.0118

[39] Kumar NH, Kluever V, Barth E, et al. (2024) Comprehensive transcriptome analysis reveals altered mRNA splicing and post-transcriptional changes in the aged mouse brain. Nucleic Acids Res 52(6): 2865–2885. 10.1093/nar/gkae172

[40] Leroux L, Desbois P, Lamotte L, et al. (2001) Compensatory responses in mice carrying a null mutation for Ins1 or Ins2. Diabetes 50 Suppl 1: S150–153. 10.2337/diabetes.50.2007.s150

[41] D’Hooge R, De Deyn PP (2001) Applications of the Morris water maze in the study of learning and memory. Brain Res Brain Res Rev 36(1): 60–90. 10.1016/s0165-0173(01)00067-4

[42] Bannerman DM, Grubb M, Deacon RM, Yee BK, Feldon J, Rawlins JN (2003) Ventral hippocampal lesions affect anxiety but not spatial learning. Behav Brain Res 139(1-2): 197–213. 10.1016/s0166-4328(02)00268-1

[43] Fanselow MS, Dong HW (2010) Are the dorsal and ventral hippocampus functionally distinct structures? Neuron 65(1): 7–19. 10.1016/j.neuron.2009.11.031

[44] Liu Z, Wan X, Chen J, et al. (2025) Genomic locus of lncRNA-Gm26793 forms an inter-chromosomal interaction with Cubn to ensure proper stem cell differentiation in vitro and in vivo. Cell Discov 11(1): 53. 10.1038/s41421-025-00805-0

[45] Milstein JL, Ferris HA (2021) The brain as an insulin-sensitive metabolic organ. Mol Metab 52: 101234. 10.1016/j.molmet.2021.101234

[46] Szabat M, Page MM, Panzhinskiy E, et al. (2016) Reduced Insulin Production Relieves Endoplasmic Reticulum Stress and Induces beta Cell Proliferation. Cell Metab 23(1): 179–193. 10.1016/j.cmet.2015.10.016

[47] Neu SC, Pa J, Kukull W, et al. (2017) Apolipoprotein E genotype and sex risk factors for Alzheimer disease: a meta-analysis. JAMA neurology 74(10): 1178–1189

[48] Fleischer AW, Frick KM (2023) New perspectives on sex differences in learning and memory. Trends in Endocrinology & Metabolism 34(9): 526–538. 10.1016/j.tem.2023.06.003

[49] Grünblatt E, Salkovic-Petrisic M, Osmanovic J, Riederer P, Hoyer S (2007) Brain insulin system dysfunction in streptozotocin intracerebroventricularly treated rats generates hyperphosphorylated tau protein. Journal of neurochemistry 101(3): 757–770

[50] Plaschke K, Kopitz J, Siegelin M, et al. (2010) Insulin-resistant brain state after intracerebroventricular streptozotocin injection exacerbates Alzheimer-like changes in Tg2576 AβPP-overexpressing mice. Journal of Alzheimer’s disease 19(2): 691–704

[51] Lester-Coll N, Rivera EJ, Soscia SJ, Doiron K, Wands JR, de la Monte SM (2006) Intracerebral streptozotocin model of type 3 diabetes: relevance to sporadic Alzheimer’s disease. Journal of Alzheimer’s Disease 9(1): 13–33

[52] Chen W, Cai W, Hoover B, Kahn CR (2022) Insulin action in the brain: cell types, circuits, and diseases. Trends Neurosci 45(5): 384–400. 10.1016/j.tins.2022.03.001

[53] Dixon-Salazar TJ, Fourgeaud L, Tyler CM, Poole JR, Park JJ, Boulanger LM (2014) MHC class I limits hippocampal synapse density by inhibiting neuronal insulin receptor signaling. J Neurosci 34(35): 11844–11856. 10.1523/jneurosci.4642-12.2014

[54] Soto M, Cai W, Konishi M, Kahn CR (2019) Insulin signaling in the hippocampus and amygdala regulates metabolism and neurobehavior. Proceedings of the National Academy of Sciences 116(13): 6379–6384. doi:10.1073/pnas.1817391116

[55] Cai W, Xue C, Sakaguchi M, et al. (2018) Insulin regulates astrocyte gliotransmission and modulates behavior. J Clin Invest 128(7): 2914–2926. 10.1172/jci99366

[56] Taib B, Deme P, Gupta S, et al. (2025) Insulin acts on astrocytes to shift their substrate preference to fatty acids. iScience 28(4): 111642. 10.1016/j.isci.2024.111642

[57] Fernandez AM, Martinez-Rachadell L, Navarrete M, et al. (2022) Insulin regulates neurovascular coupling through astrocytes. Proc Natl Acad Sci U S A 119(29): e2204527119. 10.1073/pnas.2204527119

[58] Leclerc M, Bourassa P, Tremblay C, et al. (2022) Cerebrovascular insulin receptors are defective in Alzheimer’s disease. Brain 146(1): 75–90. 10.1093/brain/awac309

