## Supplementary material for "Loss of brain insulin production impairs learning and memory in female mice": ESM

**Electronic supplementary material (ESM):**

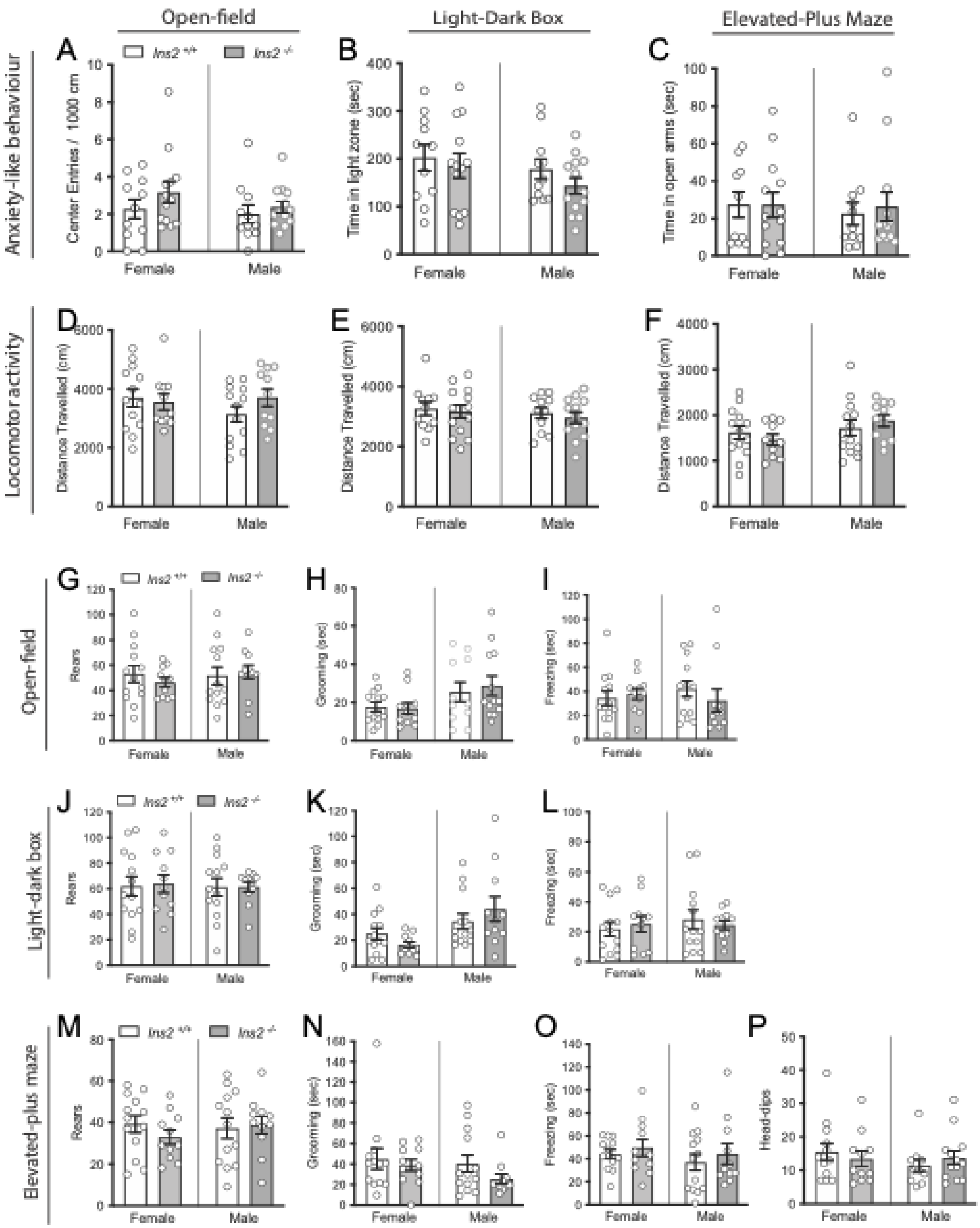


**ESM Figure 1. *Ins2*$^{-/-}$ and *Ins2*$^{+/+}$ mice do not differ in anxiety-like behaviour or locomotor activity**. **(A-C)** Anxiety-like behaviour in male and female, *Ins2*$^{-/-}$ and *Ins2*$^{+/+}$ mice (time in anxiogenic regions) on each of the open-field, light-dark box and elevated-plus maze. **(D-F)** Locomotor activity in male and female, *Ins2*$^{-/-}$ and *Ins2*$^{+/+}$ mice (time in anxiogenic regions) on each of the open-field, light-dark box and elevated-plus maze. **(G-I)** Measures of species-typical behaviours (rears, grooming, freezing) in the open-field. **(J-L)** Measures of species-typical behaviours (rears, grooming, freezing) in the light-dark box. **(M-P)** Measures of species-typical behaviours (rears, grooming, freezing, open-arm, head-dips) in the elevated plus maze. Mice were 4-5 months of age.

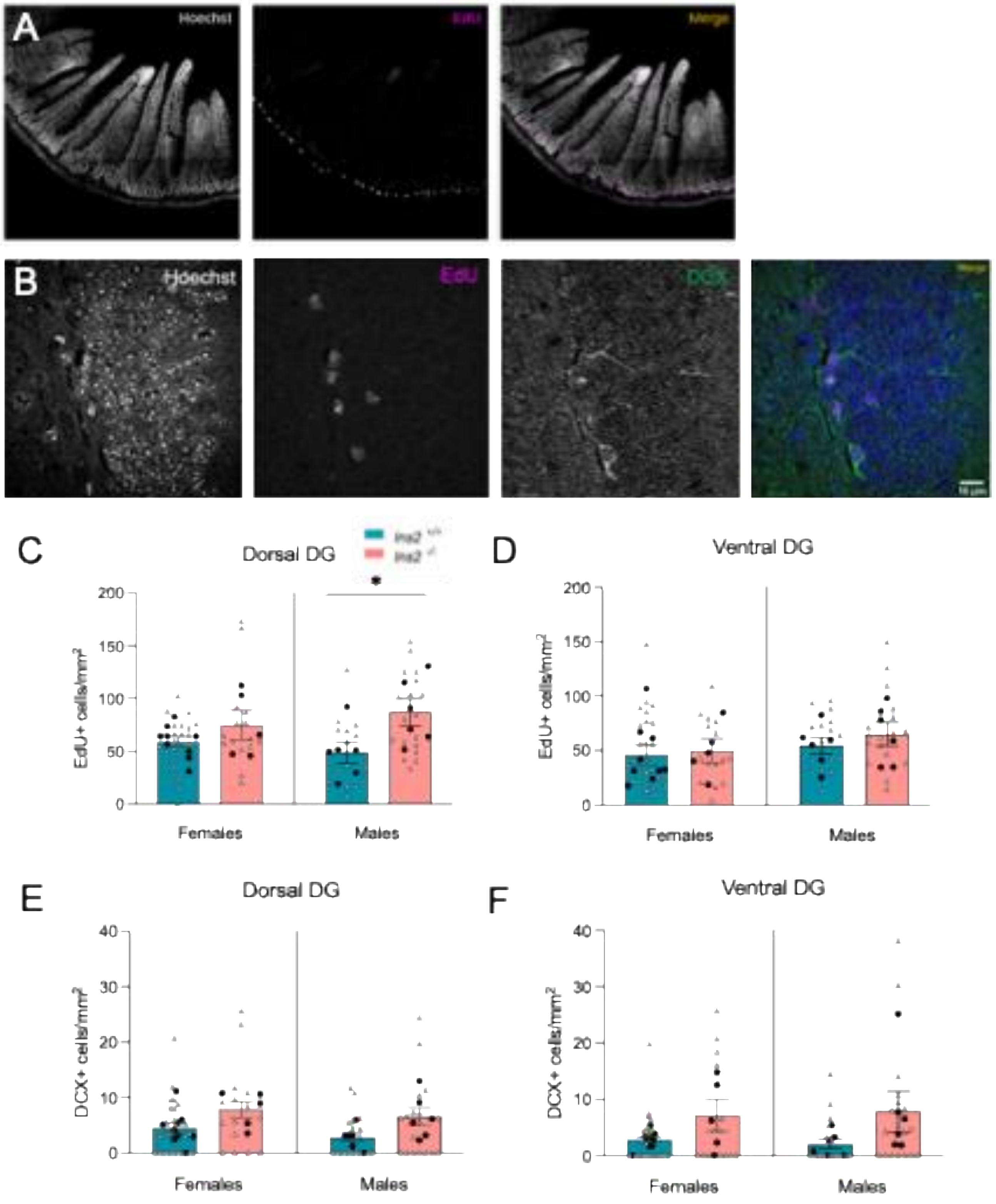


**ESM Figure 2. Effects of *Ins2* loss on adult hippocampal neurogenesis in aged male and female mice. (A)** Specific labeling of EdU in the intestine, our positive control for proliferating cells in aged animals. **(B)** Co-labeling of EdU and DCX markers in the dentate gyrus of the hippocampus. Scale bar, 10um **(C,D)** Quantification of the density of EdU$^+$ cells in the dorsal and ventral dentate gyrus in $Ins2^{-/-}$ and $Ins2^{+/+}$ mice. **(E,F)** Quantification of the density of DCX$^+$ cells in the dorsal and ventral dentate gyrus.*=p<.05. Mice were 20 months of age.
